# Paternal knockdown of tRNA (cytosine-5-)-methyltransferase (*Dnmt2*) increases offspring susceptibility to infection in red flour beetles

**DOI:** 10.1101/422063

**Authors:** Nora K E Schulz, Fakry F. Mohamed, Lai Ka Lo, Robert Peuß, Maike F de Buhr, Joachim Kurtz

## Abstract

Intergenerational effects from fathers to offspring are increasingly reported from diverse organisms, but the underlying mechanisms remain speculative. Paternal trans-generational immune priming (TGIP) was demonstrated in the red flour beetle *Tribolium castaneum*: non-infectious bacterial exposure of fathers protects their offspring against an infectious challenge for at least two generations. Epigenetic processes, such as cytosine methylation of nucleic acids, have been proposed to enable transfer of information from fathers to offspring. Here we studied a potential role in TGIP of the *Dnmt2* gene (renamed as *Trdmt1* in humans), which encodes a highly conserved enzyme that methylates different RNAs, including specific cytosines of a set of tRNAs. *Dnmt2* has previously been reported to be involved in intergenerational epigenetic inheritance in mice and protection against viruses in fruit flies. We first studied gene expression and found that *Dnmt2* is expressed in various life history stages and tissues of *T. castaneum*, with high expression in the reproductive organs. RNAi-mediated knockdown of *Dnmt2* in fathers was systemic, slowed down offspring larval development and increased mortality of the adult offspring upon bacterial infection. However, these effects were independent of bacterial exposure of the fathers. In conclusion, our results point towards a role of *Dnmt2* for paternal effects, while elucidation of the mechanisms behind paternal TGIP needs further studies.

## Introduction

Phenotypic plasticity is often enabled by epigenetic mechanisms (Duncan et al., 2014; Feinberg, 2007). An important epigenetic modification of nucleic acids is the covalent binding of a methyl group to a cytosine followed by a guanine, i.e., CpG methylation (Lyko, 2018). In insects, CpG methylation of DNA has been extensively studied (Bewick et al., 2017; Provataris et al., 2018) and is involved in many processes, such as caste determination and phase polyphenism (Elango et al., 2009; Ernst et al., 2015; Falckenhayn et al., 2013; Kucharski et al., 2008; Pasquier et al., 2014). However, this cytosine methylation not only occurs on DNA, but also on a variety of RNAs (Rana and Ankri, 2016). Cytosine methylation is facilitated by a conserved family of enzymes called DNA methyltransferases (Dnmts), which are found in most but not all animals (Goll and Bestor, 2005; Lyko, 2018). *Dnmt2* is the evolutionary most conserved member of this gene family. It can be found in many fungi, plant, and animal species, sometimes occurring in the absence of any other functional DNA methylation machinery (Durdevic and Schaefer, 2013). While *Dnmt1* and *Dnmt3* are responsible for modifications of DNA, *Dnmt2* does not methylate DNA, but rather three types of tRNA at the C38 position (Goll et al., 2006; Raddatz et al., 2013; Schaefer et al., 2010). Because of this function, the human *Dnmt2* gene received a new name: *Trdmt1* (tRNA aspartic acid methyltransferase 1). However, the gene is phylogenetically related to *Dnmt* genes, and both names remain in use (Jeltsch et al., 2017; Lyko, 2018). The methylation mark provided by *Dnmt2* protects the tRNA molecule against cleavage, which can be induced by different stressors (Schaefer et al., 2010). It has been shown that tRNA-derived small RNAs (tsRNAs) regulate mRNAs and therefore differences in tRNA cleavage could lead to altered phenotypes. In mice, dietary stress can cause increased fragmentation of tRNAs, and the altered levels of tsRNAs can lead to the transmission of the resulting metabolic phenotype from father to offspring (Chen et al., 2016; Sharma et al., 2016). This paternal transmission is dependent on *Dnmt2*, which demonstrates the importance of this gene in non-genetic inheritance (Zhang et al., 2019, 2018). Furthermore, *Dnmt2* is essential to the non-Mendelian inheritance of two paramutations in mice (Kiani et al., 2013). Both epigenetic modulations - one in the *Kit* gene leading to changes in fur color, the other in the *Sox9* gene causing overgrowth - depend on DNMT2 to be transmitted to the offspring generation (Kiani et al., 2013). The function of *Dnmt2* has also been studied in *Drosophila melanogaster*. Mutants lacking *Dnmt2* were less protected against a variety of stressors (Genenncher et al., 2018; Schaefer et al., 2010). Additionally, other studies in insects have demonstrated that *Dnmt2* plays a crucial role in managing endogenous and exogenous RNA stress and defense against RNA viruses (Bhattacharya et al., 2021a, 2017; Durdevic et al., 2013a; Phalke et al., 2009). Thus, the picture emerges that *Dnmt2* is involved in immune responses and aids in defending against or adapting to viruses and potentially other pathogens (Durdevic and Schaefer, 2013).

Phenotypic plasticity is widespread in immune defense, both within and across generations. A wealth of studies in invertebrates now provides evidence for immune priming, i.e., enhanced survival upon a secondary encounter with a pathogen (Milutinović and Kurtz, 2016; Roth et al., 2018; Schmid-Hempel, 2005). While maternal transfer of immunity appears to be a relatively common phenomenon and recent studies are beginning to shed some light on how it functions (Mondotte et al., 2020; Tetreau et al., 2019), reports about paternal transgenerational immune priming (TGIP) are scarce (Eggert et al., 2014; Roth et al., 2010) and its underlying mechanisms remain elusive. However, one thing we know about paternally transferred immune priming is that it can be not only an intergenerational but rather a transgenerational phenomenon, i.e., the protective effect of the immune priming is passed on to more than one offspring generation (Schulz et al., 2019). The paternal route of priming narrows down the possibilities by which effectors or information could be transferred from father to offspring and thus makes the involvement of epigenetic modifications, such as methylation of sperm RNA, especially likely (Roth et al., 2018). Finally, in another beetle, *Tenebrio molitor* priming of adults and larvae decreased overall RNA methylation within the generation, hinting at a possible involvement of RNA-methylating processes (Castro-Vargas et al., 2017).

*T. castaneum* possesses two *Dnmt* genes: one *Dnmt1* and one *Dnmt2/Trdmt1* homolog (Richards et al., 2008). The beetle seems to lack any functional levels of CpG DNA methylation (Bewick et al., 2017; Schulz et al., 2018; Zemach et al., 2010), but *Dnmt1* is nevertheless expressed across all life stages and is needed for proper embryonic development (Schulz et al., 2018). To our knowledge no research has been dedicated yet to study the role and function of *Dnmt2* in *T. castaneum*. We therefore used gene expression analysis via RT qPCR and systemic RNAi knockdowns to investigate the role of this gene for paternal intergenerational effects. We combined a paternal knockdown of *Dnmt2* with a non-infectious bacterial exposure treatment. We monitored offspring development and survival of bacterial infection, to investigate whether *Dnmt2* is involved in and possibly provides the epigenetic mechanism behind paternal immune priming.

## Results

### Expression of *Dnmt2*

Before investigating a possible function of *Dnmt2* in *T. castaneum*, we monitored expression of this gene throughout the life cycle of the beetle (i.e., in eggs, larvae, pupae and adults). The levels of *Dnmt2* transcripts (relative to two housekeeping genes) in eggs and pupae did not differ from adults (eggs: relative expression=0.932, n=4 pools of 500-1000 eggs, p=0.76; pupae: relative expression=0.989, n=8 pools of 6 individuals, p=0.94; relative to virgin adults, n=8 pools of 6 individuals; Figure 1A). Larvae expressed *Dnmt2*, but significantly less than adults (relative expression=0.352, n=7 pools of 10 individuals, p<0.001; Figure 1A). Additionally, *Dnmt2* expression did not differ between males and females in neither the pupal (female: relative expression=0.784, n=4 pools of 6 individuals, p=0.23) nor the adult stage (female: relative expression=0.709, n=4 pools of 6 individuals, p=0.14).

**Figure 1.**
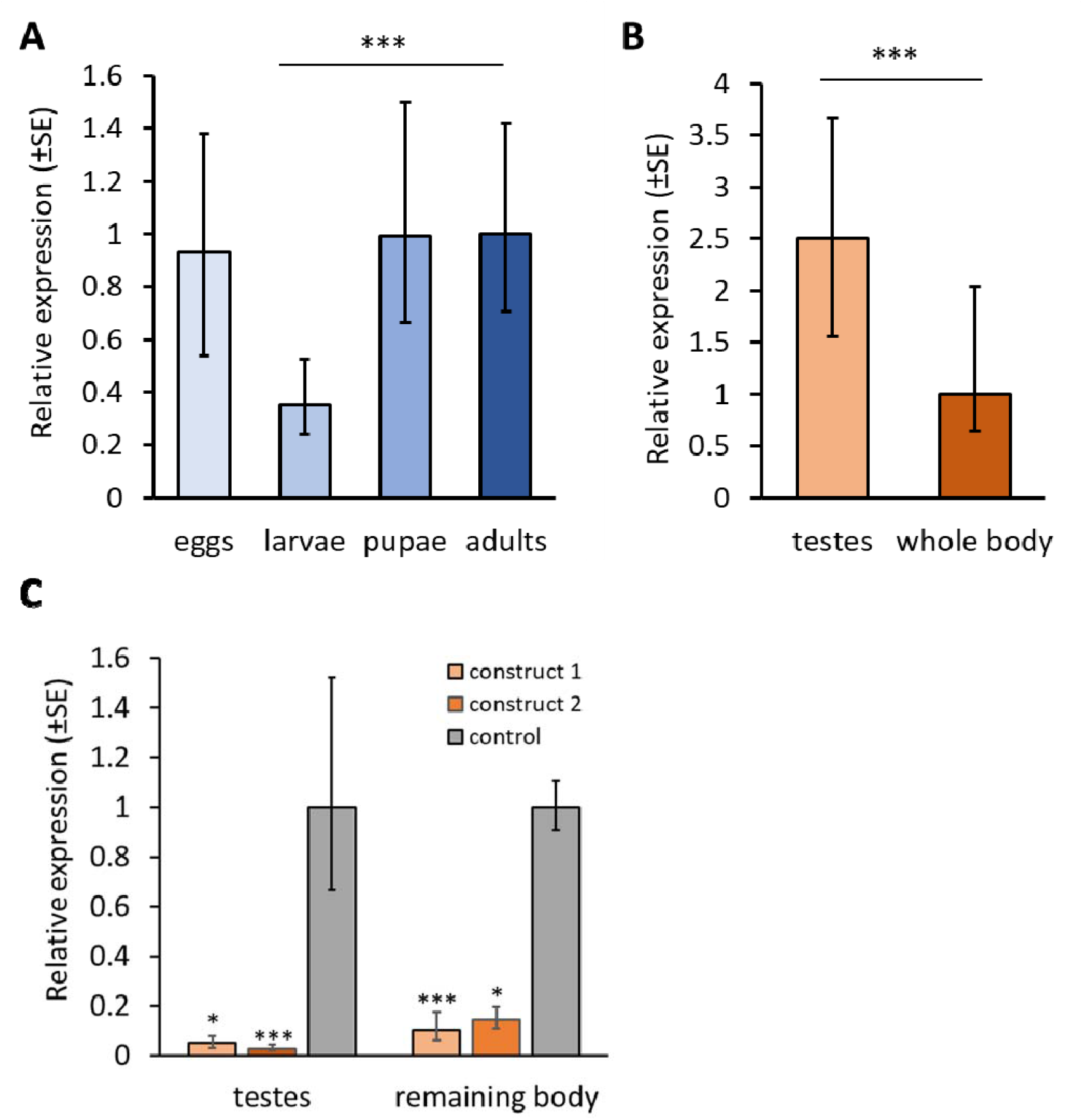
*Dnmt2* gene expression and knockdown. **A)** Relative expression in four distinct life stages of *T. castaneum* compared to adult samples (eggs: n=4 pools of 500-1000 eggs, 24h-48h post oviposition; larvae: n=7 pools of 10 larvae, 14-19 dpo; pupae: n=8 pools of 6 individuals; adults: n=8 pools of 6 individuals, one week after eclosion). **B)** Relative expression in testes and male whole-body samples (n=6 pools of 8-10 individuals). **C)** *Dnmt2* knockdown in testes and remaining body parts. Relative expression of *Dnmt2* after *Dnmt2* dsRNA injection of one of two constructs compared to samples from the same tissue of the RNAi control treatment (GFP). Tissue from 6-8 individuals was pooled per sample (GFP-dsRNA n=3, Dnmt2-dsRNA construct 1 n=4, construct2 n=4). Asterisks indicate significant differences (* =p<0.05, ***= p<0.001) according to statistical analysis using Wise Fixed Reallocation Randomization Test (REST2009 software; Pfaffl et al., 2002).

Furthermore, we analyzed the expression of *Dnmt2* in the reproductive tissue of the male beetles and compared it to whole body samples of the same sex, because expression in the testes could hint at a possible relevance of *Dnmt2* in male reproduction or even an involvement in the transfer of information from father to offspring as possibly needed for TGIP. *Dnmt2* mRNA levels in the testes were significantly higher than in whole-body samples (relative expression=2.497, n=6 pools from 8 individuals, p=0.001; Figure 1B).

### Strong systemic *Dnmt2* knockdown including the testes

We used two non-overlapping dsRNA constructs to knockdown *Dnmt2* in pupae and tested whether the downregulation extends to the gonads in the adult males. With both constructs, we saw significantly lowered expression of *Dnmt2* in the testes as well as the remaining bodies of the same beetles (Figure 1C). In the testes samples, expression of *Dnmt2* was significantly downregulated by 95.0% and 97.2% in construct 1 (n=4 pools of 4-8 individuals, p=0.019) and construct 2 (n=4 pools of 4-8 individuals, p<0.001), respectively, compared to the RNAi control (n=3). The effect was similar for the remaining body tissue, where expression was down by 89.6% (p=0.001) for construct 1 and 85.7% (p=0.024) for construct 2, compared to the RNAi control. This confirms that pupal dsRNA injection causes a systemic knockdown of *Dnmt2*, which includes the male gonads.

### Paternal *Dnmt2* knockdown and TGIP

To determine whether *Dnmt2* is involved in the paternal transfer of immunity, we combined the *Dnmt2* knockdown with the injection of heat-killed bacteria into the fathers, and exposed the offspring to a bacterial challenge, so as to test for paternal TGIP (Figure 2).

**Figure 2.**
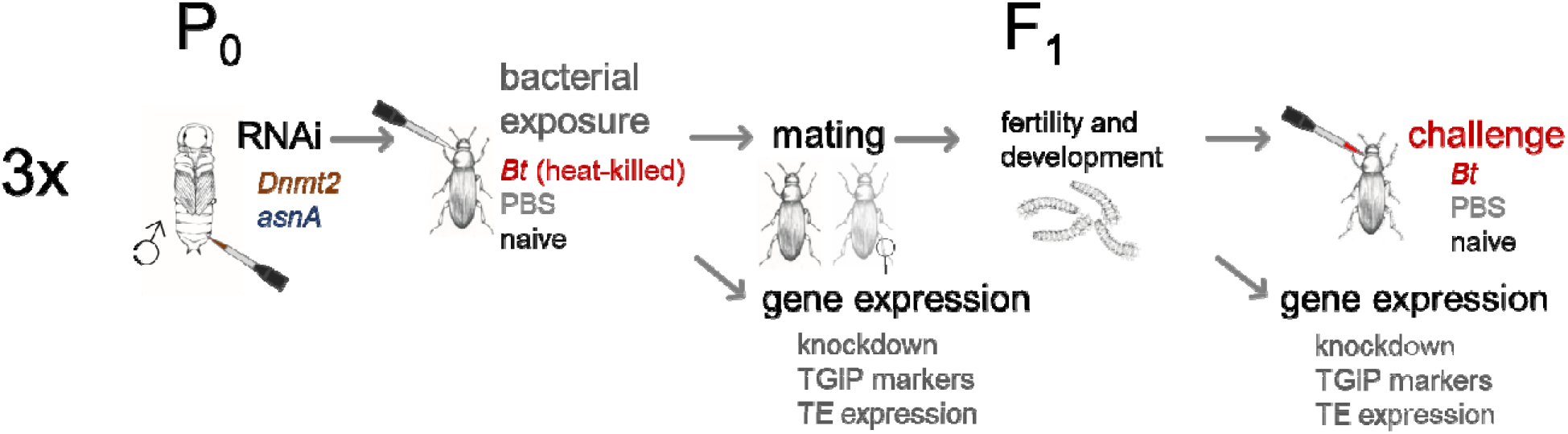
Overview of the experimental design combining paternal RNAi-mediated knockdown of *Dnmt2* with paternal TGIP. The entire experiment was conducted in three replicates (*asnA*=RNAi control; PBS=injection control; naïve=handling control).

### Survival of RNAi and bacterial injections

The RNAi injections with *Dnmt2* or asnA control dsRNA did not increase mortality, neither directly following the treatment of the pupae (GLMM, df=1, *X* ^2^=0.04, p=0.84, Figure S1a), nor after the injection of heat-killed bacteria in the mature adults ten days later (GLMM, df=1, *X* ^2^=0.16, p=0.69; Figure S1b). The latter injection itself slightly increased mortality, for both heat-killed bacteria and sterile PBS controls, which we attribute to the wounding during these injections, as none of the naïve controls died (GLMM, df=2, *X* ^2^=15.89, p<0.001; Figure S1b).

### Paternal knockdown of *Dnmt2* does not persist into the offspring generation

We tested for persistence of the knockdown of *Dnmt2* after pupal RNAi in a subgroup of the adults, one day after the bacterial injection, i.e., 13 days after knockdown. *Dnmt2* was still weakly, but significantly downregulated (to between 8.8% and 17.7%) compared to RNAi control, regardless of the received bacterial injection treatment (**Error! Reference source not found**.A). As expected, *Dnmt2* mRNA had returned to normal levels in the adult offspring and there were no significant differences between the RNAi treatments (**Error! Reference source not found**.B). Additionally, the paternal bacterial injections did not affect *Dnmt2* expression in the adult offspring (**Error! Reference source not found**.B).

### Knockdown of *Dnmt2* and bacterial injection did not affect male fertility

Neither the knockdown of *Dnmt2* nor the injection of heat-killed bacteria appear to affect the fitness of the treated individuals, as neither treatment significantly altered male fertility. The number of live offspring obtained from a 24 h single pair mating period did not differ significantly for either of the treatments (GLMM: RNAi, df=1, *X* ^2^=2.11, p=0.15; bacterial injection, df=2, *X* ^2^=0.44, p=0.80; Figure S2).

### Paternal knockdown of *Dnmt2* but not bacterial injection affected offspring development

We monitored offspring development by measuring the proportion of pupae over three consecutive days and the proportion of eclosed adults 26 days post oviposition (dpo). Animals from all six treatment combinations of RNAi and bacterial injection showed similar pupation rates 21 and 22 dpo (Figure 3C, Figure S3a). However, at 23 dpo significantly less larvae had reached pupation in the *Dnmt2* paternal knockdown group than in the RNAi control, independent of paternal bacterial injection treatment (GLMM: RNAi, df=1, *X* ^2^=3.9, p<0.05; bacterial injection, df=2, *X* ^2^=0.19, p=0.91; Figure 3C). The proportion of eclosed adults 26 dpo was not significantly affected by any paternal treatment (GLMM: RNAi, df=1, *X* ^2^=0.04, p=0.84; heat-killed bacterial exposure, df=2, *X* ^2^=0.48, p=0.79; Figure S3b).

**Figure 3.**
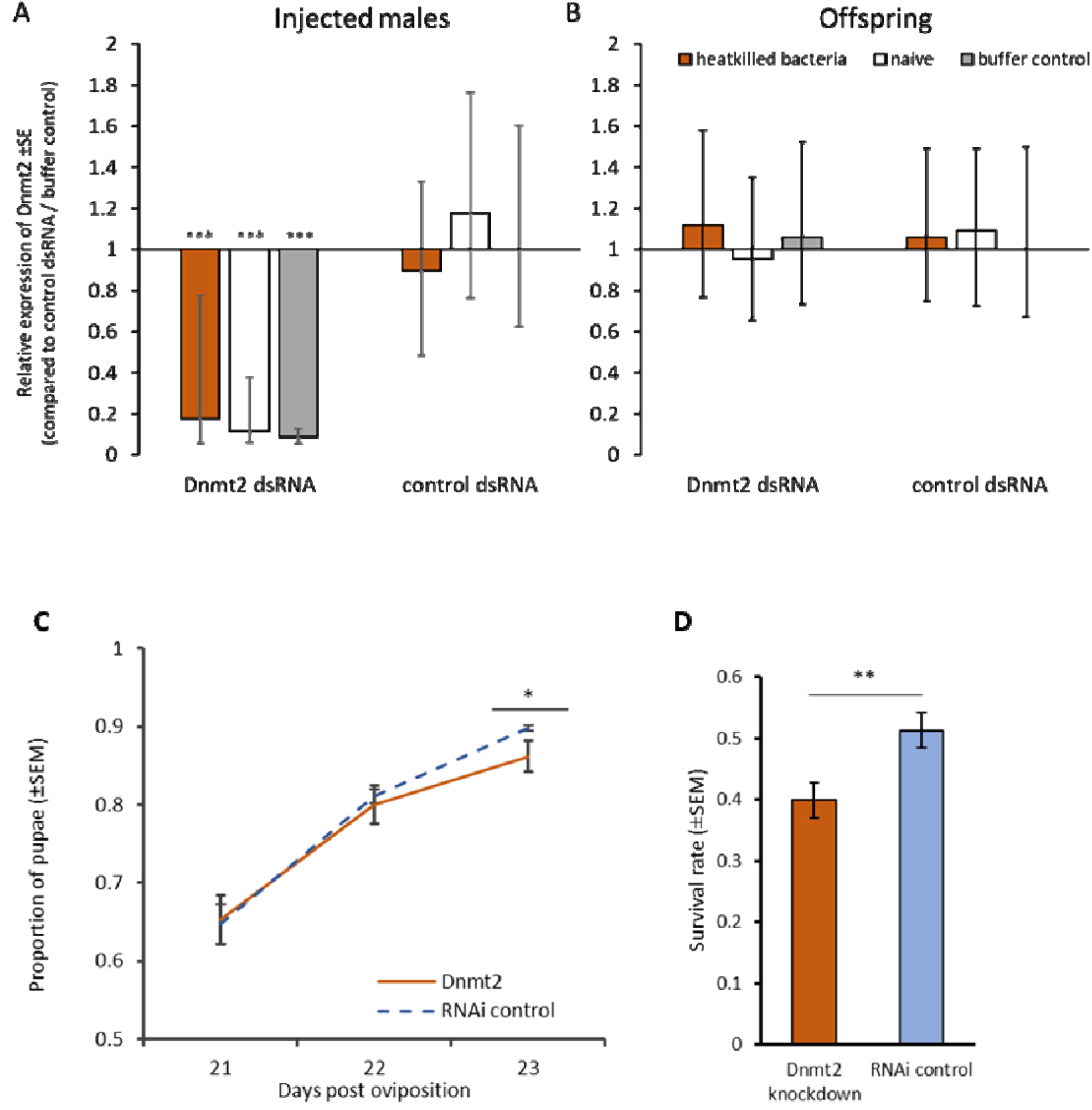
Paternal knockdown of *Dnmt2* and its effects on the offspring. **A)** Relative *Dnmt2* expression in males of the parental generation for the three heat-killed bacterial exposure treatments (heat killed bacteria, naïve and buffer control) and two RNAi treatments (*Dnmt2* dsRNA and control dsRNA). Each group was compared against the control dsRNA/ buffer control group (n=12-14). Asterisks indicate significant differences (p<0.001) according to statistical analysis using Pair Wise Fixed Reallocation Randomization Test (REST2009 software, Pfaffl et al., 2002). **B)** Relative *Dnmt2* expression in adults of the F1 generation for the three paternal heat-killed bacterial exposure treatments (heat killed bacteria, naïve and buffer control) and two RNAi treatments (*Dnmt2* dsRNA and control dsRNA). Each group was compared against the control dsRNA/ buffer control group (n=12-15). **C)** Pupation rate of F_1_ generation 21-23 dpo (±SEM for three experimental replicates) **D)** Survival of the F_1_ generation after bacterial challenge according to paternal RNAi treatment. Shown are the proportions of adults that were alive four days post injection with *B. thuringiensis* (±SEM for three experimental replicates). Asterisks indicate significant differences (*=p<0.05, **=p<0.01).

### Paternal *Dnmt2* knockdown reduces offspring survival after bacterial challenge

Finally, we injected adult beetles from the offspring generation with a potentially lethal dose of *B. thuringiensis* to test for paternal trans-generational immune priming and whether this was affected by the downregulation of *Dnmt2* in the fathers. In this experiment, injection of heat-killed bacteria into fathers did not affect offspring survival after bacterial challenge (GLMM, df=2, *X* ^2^= 0.17, p=0.92; Figure S4), which can possibly be explained by the unavoidable additional wounding that all fathers were subjected to because of the RNAi injection treatment. However, offspring of individuals that had received *Dnmt2* knockdown were significantly less likely to survive the bacterial challenge (GLMM, df=2, *X* ^2^=7.78, p<0.01; Figure 3D), demonstrating that *Dnmt2* is involved in paternal effects such that knock-down increases the offspring’s susceptibility to pathogens in the beetle.

### Expression of TGIP marker genes and TEs is not affected by *Dnmt2* knockdown

In fathers and offspring alike, we measured the expression of three genes, which are related to stress or immune responses and were previously shown to be upregulated in the adult offspring of fathers, injected with heat-killed bacteria (Eggert et al., 2014). By measuring the expression in the fathers, we intended to see whether these genes would already be affected within the treated generation. None of the three candidate genes (*hsp83, nimB* and *PGRP*) showed any significant differential expression neither in the paternal nor in the adult offspring generation (Table S2).

For the same animals from the paternal generation, we also measured the expression of seven TEs. Genenncher *et al*. (2018) observed that the absence of *Dnmt2* and the exposure to heat stress led to the activation and accumulation of certain TEs in *D. melanogaster*. Here, we could not observe any significant upregulation of the expression of TEs after wounding or bacterial injection in the knockdown or control treatment (Table S3).

## Discussion

*Dnmt2* can be found in almost all eukaryote species and is the most conserved member of the *Dnmt* family (Schaefer and Lyko, 2010). It also has a function in some organisms lacking one or both other *Dnmt* genes and which are often devoid of any functional DNA methylation system (Durdevic and Schaefer, 2013). This also appears to be the case in *T. castaneum*, which lacks *Dnmt3* and does not have any functional CpG DNA methylation, but still expresses *Dnmt2* (Bewick et al., 2017; Richards et al., 2008; Schulz et al., 2018; Zemach et al., 2010). Although, we know today that *Dnmt2* methylates tRNA and not DNA, it remains unclear what the role of this epigenetic mechanism is. We observed that *Dnmt2* mRNA transcripts are present at relatively low but consistent levels in all life stages and in both sexes of the beetle, therefore the enzyme likely functions in both males and females and throughout the entire life cycle. In fruit flies, mice, and humans, *Dnmt2* or its mammalian ortholog *Trdmt1* canonically methylate a small set of tRNAs (Goll et al., 2006; Jurkowski et al., 2008; Schaefer et al., 2010), which are highly abundant in sperm (Peng et al., 2012) and have been shown to be involved in paternal transmission of metabolic phenotypes in mice (Chen et al., 2016; Sharma et al., 2016). The significantly higher expression of *Dnmt2* that we observed in testes of *T. castaneum* could hint at the involvement of *Dnmt2* in paternal epigenetic inheritance also in this beetle.

We used systemic paternal RNAi, to investigate how a *Dnmt2* knockdown would affect offspring phenotypes and to discover a potential role of this gene in intergenerational epigenetic inheritance, specifically from father to offspring. We were able to show that the *Dnmt2* knockdown extended to the testes, where *Dnmt2* transcripts were strongly reduced, which is a likely prerequisite for manipulating the transfer of information between fathers and offspring. The offspring of *Dnmt2* RNAi-treated fathers needed longer to reach pupation and thus remained longer in the vulnerable larval stage, e.g., the stage susceptible to oral bacterial infection (Milutinović et al., 2015; Schulz et al., 2019). They also died at a higher rate from a *B. thuringiensis* infection, which may point towards a generally higher stress sensitivity. In recent years, it has become clear that biological functions of *Dnmt2* are more easily detected under stress conditions (Durdevic and Schaefer, 2013). Increased sensitivity to thermal and oxidative stress has been observed in *D. melanogaster Dnmt2* mutants (Schaefer et al., 2010), while overexpression of the same gene led to increased stress tolerance (Lin et al., 2005). During the stress response, *Dnmt2* appears to control the fragmentation of tRNA and can be located at cellular stress compartments (Durdevic et al., 2013b; Schaefer et al., 2010). More recently, it has also been demonstrated that the knockout of *Dnmt2* leads to a faster decline of immune function with age in adult flies (Abhyankar et al., 2018). Also in *Drosophila, Dnmt2* deficient mutants are not capable of efficient pathogen blocking and carry a higher viral load (Bhattacharya et al., 2017). Finally, its absence disrupts the small interfering RNA pathway by inhibiting dsRNA degradation by *Dicer* (Durdevic et al., 2013b). However, all these effects occur in the treated generation and have not been tested in the offspring. In our case the developmental and immunological effects were observed in the offspring of treated fathers, even though offspring themselves exhibited normal *Dnmt2* expression, which suggests intergenerational effects. A role in intergenerational paternal effects for *Dnmt2* has so far only been established in mice, where the gene is essential for the transmission of an acquired metabolic disorder (Zhang et al., 2018). Therefore, further studies are needed to clarify the role that *Dnmt2* plays in enabling appropriate stress responses within and across generations.

Finally, we combined the knockdown of *Dnmt2* with a paternal heat-killed bacterial exposure treatment, to determine whether the activity of the tRNA (cytosine-5-)-methyltransferase affects the transfer of information on bacterial encounter from father to offspring. We did not observe any paternal immune priming in the present study, as offspring survival of bacterial infection was independent of the paternal heat-killed bacterial exposure treatment. Furthermore, we did not observe any upregulation in the previously described marker genes for immune and stress responses (Eggert et al., 2014) nor any changes in the expression of transposable elements. This surprising lack of any priming effect (in contrast to previous studies: (Eggert et al., 2014; Roth et al., 2010; Schulz et al., 2019) might have been caused by the wounding associated with the injection of dsRNA into all fathers for the pupal RNAi treatment, which took place before the injection of the heat-killed bacteria (note that we included an RNAi control, but no fully naïve control for the RNAi treatment, Figure 2 Overview of the experimental design combining paternal RNAi-mediated knockdown of *Dnmt2* with paternal TGIP. The entire experiment was conducted in three replicates (*asnA*=RNAi control; PBS=injection control; naïve=handling control).). The wounding associated with the dsRNA injection might have activated immune responses, such that all animals could already have entered a ‘primed’ state. To our knowledge there are no studies that directly address the question how injuries during the pupal phase influence later immune responses. However, some experiments show that wounding in control treatments also to a certain extent may increase survival of a subsequent bacterial challenge (Roth et al., 2010; Tate et al., 2017). Alternatively, the pupal RNAi injections might have inhibited effective immune priming. Lastly, although TGIP in *T. castaneum* is robust and repeatable (Eggert et al., 2014; Roth et al., 2010; Tate et al., 2017), it also has become apparent that this phenomenon cannot be observed in every experiment (Tate et al., 2017) and beetle population (Khan et al., 2016).

We did not observe any effects of the *Dnmt2* knockdown on expression of the studied TEs, in contrast to *D. melanogaster Dnmt2* mutants that showed increased TE expression (Genenncher et al., 2018). However, further studies are needed to make any firm conclusions, because the lack of expression differences for a limited set of TEs does not exclude the possibility that *Dnmt2* plays a role in the regulation of TEs in *T. castaneum*. In plants, flies, and mice the absence of *Dnmt2*/*Trdmt1* is not lethal under standard conditions and mutants remain fertile (Goll et al., 2006). The same appears to be true in the case of *T. castaneum*, where we did not observe any mortality nor apparent phenotypic changes after a significant downregulation of *Dnmt2*. Additionally, male fertility was not affected by the knockdown under *ad libitum* conditions. Therefore, the maintenance of knockout lines appears feasible, which makes this gene a suitable target for CRISPR/Cas knockout to further study its function without the necessity of repeated RNAi injections for each experiment. Moreover, methylation-sensitive sequencing of sperm RNA could further elucidate the underlying molecular processes.

In conclusion, our study for the first time describes paternal effects related to *Dnmt2* in an invertebrate. The here observed prolonged development and increased susceptibility to infection occurred in the presence of normal *Dnmt2* expression in the offspring. Therefore, tRNA methylation in sperm provides a fascinating possibility for transmitting information from fathers to offspring through paternal epigenetic inheritance.

### Experimental procedures

#### Model organism

The red flour beetle, *T. castaneum* is a well-established model organism with a fully sequenced genome (Richards et al., 2008) and diverse molecular tools (Bucher et al., 2002; Gilles and Averof, 2014; Peuß et al., 2016; Schmitt-Engel et al., 2015) readily available. For this study, the *T. castaneum* line Cro 1 was used, which was established from 165 pairs of wild caught beetles collected in Croatia in June 2010 (Milutinović et al., 2013). Beetles were maintained in plastic breeding boxes with foam stoppers to ensure air circulation. Standard breeding conditions were 30°C and 70% humidity with a 12-hour light/dark cycle. As food source 250g of heat sterilized (75°C) organic wheat flour containing 5% brewer’s yeast were given.

### RT-qPCR to measure gene expression of *Dnmt2*

To assess the expression of *Dnmt2* (gene ID: LOC663081, TcasGA2_TC005511) throughout the life cycle of the beetle, the four distinct life stages were sampled: eggs (n=4 pools of 500-1000 eggs, 24h-48h post oviposition), larvae (n=7 pools of 10 larvae, 14-19 days post oviposition (dpo)) pupae (n=8 pools of 6 individuals), virgin adults (n=8 pools of 6 individuals, one week after eclosion). For pupae and adults, half of the pooled samples contained females and the other half males in order to test also for differential expression between the sexes. Furthermore, gonads were dissected from unmated adult males. All samples were shock frozen in liquid nitrogen. Total RNA was extracted, and genomic DNA digested by combining Trizol (Ambion RNA by Life Technologies GmbH, Darmstadt, Germany) and chloroform treatment with the use of the Total RNA extraction kit (Promega GmbH, Mannheim, Germany) as described in Eggert et al. (2014).

Extracted RNA was reverse transcribed to cDNA with the RevertAid First Strand cDNA kit (Thermo Fisher Scientific, Waltham, MA USA) using provided oligo-dTs. In the following RT qPCR with a Light-Cycler480 (Roche) and Kapa SYBR Fast (Kapa Biosystems, Sigma-Aldrich), each sample was used in two technical replicates. Further analysis was conducted as described in Eggert et al. (2014) and replicates were used in further analysis if the standard deviation between their crossing point values was below 0.5, otherwise the reaction was repeated. Previously, high primer efficiency had been confirmed and where possible it was made sure that primers crossed exon-intron boundaries (Table S1). The housekeeping genes ribosomal proteins *rp49* and *rpl13a* were used for normalization of the expression of the target genes.

### Systemic RNAi-mediated *Dnmt2* knockdown

If a knockdown of *Dnmt2* disrupts the proper transfer of paternal effects to the offspring generation, we should be able to observe a downregulation that extends to the reproductive tissues, i.e., the testes. To investigate this, we performed pupal RNAi, dissected the testes of eclosed males and compared *Dnmt2* expression in testes and remaining bodies.

For injections of dsRNA, male pupae (22 dpo) were glued with the hindmost segment of the abdomen to a glass slide to immobilize them. One glass slide held between 16 and 20 pupae. Pupae were either injected with dsRNA from one of two constructs for the target gene *Dnmt2* or, as a control for the treatment procedure, with dsRNA from a *GFP* gene, which bears no sequence similarity to any known *T. castaneum* gene (Schmitt-Engel et al., 2015). The *Dnmt2* dsRNA constructs have been previously used in the ibeetle RNAi screen (Schmitt-Engel et al., 2015)http://ibeetle-base.uni-goettingen.de/details/TC005511) and were obtained from EupheriaBiotech (Dresden, Germany). Injections were carried out with a microliter injector (FemtoJet, Eppendorf AG, Hamburg, Germany) and borosilicate glass capillaries (100 mm length, 1.0 mm outside diameter, 0.021 mm wall thickness; Hilgenberg GmbH, Malsfeld, Deutschland) using dsRNA at a concentration of 1000 ng/µl dissolved in phosphate buffered saline (PBS). We injected pupae between the second and third last segment of their abdomen. We placed glass slides with the pupae into petri dishes with flour and yeast in order to provide diet to eclosed adults.

Seven days post eclosion, we dissected the testes from the adults and collected gonad tissue and remaining bodies separately. We pooled samples from four to eight individuals to create biological replicates (*Dnmt2* construct 1 n=4, construct 2 n=4, *GFP* control n=3). RNA extractions and qPCR were performed as described above.

### Paternal *Dnmt2* knockdown and TGIP

We aimed to downregulate *Dnmt2* through paternal RNAi and to investigate whether this knockdown would affect paternal TGIP (Figure 2). For this, around 2000 adult beetles (one week old) were allowed to lay eggs for 24h. Two weeks later, larvae were collected and put into individual wells of a 96 well plate, which contained flour and yeast. The oviposition was repeated with two further, independent cohorts on the two following days, producing three experimental replicates.

### Paternal RNAi

The sex of the beetles was determined in the pupal stage, and male pupae were prepared for RNAi treatment, while females were individualized and kept for mating. We performed the RNAi treatment in a similar manner as described above with some changes. For feasibility, we conducted this part of the experiment only using dsRNA for Dnmt2 construct 1 (Table S1). We used dsRNA transcribed from the *asparagine synthetase A* (asnA) gene found in *Escherichia coli* as treatment control(Richards et al., 2008; Table S1). The dsRNA construct for the RNAi control was produced in our lab via cloning followed by PCR and *in vitro* transcription using the T7 MEGAscript Kit (Ambion by Life TechnologiesTM GmbH, Darmstadt, Gemany; Peuß et al., 2016). Finally, dsRNA was suspended in ultrapure water instead of PBS. Over the three experimental blocks a total of 583 pupae were injected with *Dnmt2* dsRNA and 585 pupae served as RNAi control and were therefore injected with *asnA* dsRNA. Eclosion and survival of the procedure were recorded daily from three to six days post injection.

### TGIP

When all surviving males from the RNAi treatment had reached sexual maturity seven days after eclosion, they were injected for priming with around 37,000 cells of heat-killed *B. thuringiensis* (DSM no. 2046, obtained from the German Collection of Microorganisms and Cell Cultures, DSMZ) suspended in PBS. This treatment has successfully been used in prior TGIP experiments (Eggert et al., 2014; Roth et al., 2010). Bacterial cultures were grown overnight as previously described (Roth and Kurtz, 2009). They were washed with PBS and heat-killed by exposure to 95°C for 30 minutes. We injected a volume of 18.4 nL of bacterial suspension containing 10^7^ cells/mL, which constitutes a LD50 dose for the beetle line (Schulz et al., 2019). Control groups were either injected with PBS (injection control) containing no bacterial cells or were left naïve. Injections were performed using the nanoliter injector Nanoject II (Drummond Scientific Company, Broomall, PA, USA) and individuals were injected between head and thorax. Beetles were kept individually before and after the injections. Survival of the bacterial injection was recorded 24h later.

### Gene expression after RNAi and heat-killed bacterial exposure treatment

Twenty-four hours post heat-killed bacterial injection, a subgroup of males was used for gene expression analysis to confirm the knockdown of *Dnmt2*. In addition to the expression of *Dnmt2*, the expression of three immunity and stress-related genes (*hsp83, nimB* and *PGRP*; Table S1) was analyzed, based on a previous study showing effects of paternal priming on gene expression (Eggert et al., 2014). For each RNAi* heat-killed bacterial injection treatment combination and block, five samples were taken consisting of a pool of two to five individuals. RNA extraction, cDNA reverse transcription and RT qPCR were performed as described above. Finally, we also analyzed the expression of seven transposable elements (TEs) (Table S1), because the absence of *Dnmt2* can cause the activation of TEs (Genenncher et al., 2018). Because of the lack of polyadenylation of some TE transcripts, we used random hexamer primers for cDNA reverse transcription in this case (Thermo Fisher Scientific, Waltham, MA USA).

### Production and development of offspring generation

One day after the heat-killed bacterial exposure, single pair matings were carried out for 24h with virgin females from the same population (n=12-50 mating pairs per treatment combination and experimental replicate). Twelve days after the oviposition to produce the F1 offspring generation, larvae from each pair were counted and up to six individuals were individualized and kept for further analyses. Additionally, one randomly chosen larva from each mating pair was used for developmental checks until it died or eclosed as an adult. The development was monitored daily from 21 to 23 dpo to check for pupation and at 26 dpo we recorded the proportion of eclosed adults.

### Gene expression in the offspring generation

One week after most of the offspring generation had eclosed, five pools per RNAi and heat-killed bacterial exposure treatment combination and experimental replicate were sampled for gene expression analysis. Each sample consisted of five adult beetles of unknown sex. To avoid pseudo-replication only one beetle per family was used. Again, the expression of *Dnmt2* and three potential TGIP marker genes (*hsp83, nimB, PGRP*) (Eggert et al., 2014) was analyzed as described above.

### Bacterial challenge of adult offspring

One week after their eclosion, adults of the F1 offspring generation were submitted to a potentially lethal bacterial injection (challenge). For this, bacteria from the same *B. thuringiensis* stock as for the paternal injections were used. Again, an overnight culture from a glycerol stock was grown in liquid medium and washed in PBS. The injection procedure was similar to the paternal exposure and again included an injection control and a naïve group. The dose was adjusted to around 370 live bacterial cells per animal. From every family one sibling each was used for the treatment and the controls. Again, beetles were kept individually before and after injection to avoid any cross contaminations. Survival of the challenge was recorded one day and four days post injection.

### Statistics

All gene expression data were analyzed with the Relative Expression Software Tool (REST2009 software; (Pfaffl et al., 2002) as described in Eggert et al. (2014) In short, this software enables comparisons between treatment and control groups of up to 16 samples, while using the ΔΔ Ct method, normalizing expression via two housekeeping genes and taking primer efficiencies into account. REST2009 uses a Pair Wise Fixed Reallocation Randomization Test to determine significant differences (Pfaffl, 2001; Pfaffl et al., 2002).

All other analyses were performed in RStudio version 0.99.467 (RStudio Team, 2015) under R version 3.3.3 (R Development Core Team, 2008) using additional packages lme4 (Bates et al., 2015) and MASS (Venables and Ripley, 2002). Survival of injections for RNAi and heat-killed bacterial exposure in the parental generation, the fertility of the treated males as well as the development of the offspring (proportion of pupae 21-23 dpo and proportion of adults 26 dpo) and their survival after bacterial challenge were analyzed in generalized linear mixed effect models (GLMMs) with the according error distributions and experimental replicate as a random factor.

## Acknowledgements

We thank Barbara Hasert and Kathrin Brüggemann for help with lab work, Jürgen Schmitz for support regarding the expression analysis of TEs and Sina Flügge for providing us with drawings of *T. castaneum*. This project was funded in part by the German Research Foundation (DFG) as part of the SFB TRR 212 (NC^3^), project number 396780003 (granted to JK) and by the Volkswagen Stiftung, project number I/84 794 (granted to MD).

## Contributions

NS, MD, and JK conceived and designed the study. NS, FM, LL, and RP conducted the experiments. NS and RP analyzed the data. NS wrote the manuscripts with comments from all authors.

## Supplementary materials

**Table S1** Primer sequences and efficiencies for RTqpCR

**Table S2** Relative expression of TGIP marker genes in parental (P_0_) and offspring (F_1_) generation compared to expression in RNAi control* bacterial exposure control group (normalized over two housekeeping genes (n=15). Statistical analysis was performed using Pair Wise Fixed Reallocation Randomization Test (REST2009 software, Pfaffl et al., 2002)

**Table S3** Relative expression of TEs after RNAi and heat-killed bacterial exposure treatment in parental (P_0_) generation (expression relative to two housekeeping genes and RNAi control* naïve group, n=6-8). Statistical analysis was performed using Pair Wise Fixed Reallocation Randomization Test (REST2009 software, Pfaffl et al., 2002)

**Figure S1** Survival of RNAi treatment a) Proportion of surviving mature adult males seven days post eclosion (n= 8-11 pools of 13-20 individuals per RNAi treatment and replicate). b) Proportion of surviving mature adults eight days post eclosion and 24 h post heat-killed bacterial exposure injection (± SEM for three experimental replicates, n=25-70 per RNAi treatment, heat-killed bacterial exposure treatment, and replicate). Different letters indicate significant differences.

**Figure S2** Male fertility after pupal RNAi treatment and adult heat-killed bacterial exposure injections. Shown is the offspring number from a 24h mating/oviposition period for three replicates (± SEM) consisting of 12-50 mating pairs each.

**Figure S3** Development of offspring after paternal heat-killed bacterial exposure with *B. thuringiensis* a) pupation rate (±SEM for three experimental replicates) b) proportion of eclosed adults (±SEM for three experimental replicates) 26 days post oviposition.

**Figure S4** Survival of F_1_ generation after bacterial challenge according to paternal heat-killed bacterial exposure treatment. Shown are the proportions of adult offspring that were alive four days post injection with live *B. thuringiensis* for three independent experimental replicates.

