## Supplemental_tables_and_figures for "Paternal knockdown of tRNA (cytosine-5-)-methyltransferase (*Dnmt2*) increases offspring susceptibility to infection in red flour beetles"

Supplementary Materials

**Table S 1** Primer sequences and efficiencies for RTqpCR

| Gene | NCBI reference | Forward primer | Reverse primer | Efficiency |
| --- | --- | --- | --- | --- |
| *Dnmt2* | XM_008198777 | ggcgttaaaagttagtggggta | ggggcgacattagaattgtg | 1.95 |
| *hsp90* | NM_001313877 | cgcagttcattggctatccc | gtcttcgccttcttcctcct | 1.97 |
| *nimB* | XM_008200605 | cacaagggaatgggaccagg | tgccattagggcagccattt | 1.94 |
| *PGRP* | XM_008194325 | ccgcgtcaaaggcattcaaa | tgccatcacccccaatcaag | 1.96 |
| *rp49* | XM_964471 | ttatggcaaactcaaacgcaac | ggtagcatgtgcttcgttttg | 1.98 |
| *rpl13a* | XM_969211 | ggccgcaagttctgtcac | ggtgaatggagccacttgtt | 2 |
| *Gypsy3* |  | gctaagcagaaacaggaaaccgat | tggtcacgtctgttaaacttagctc | 1.97 |
| *Gypsy 5* |  | cgacttttgcttcccctacctg | gtgccatctgctgaaacttcgt | 1.96 |
| *Gypsy9* |  | tttcgccgatttattgatcagttcg | tttaaagtggtgaatgcctcgtct | 1.91 |
| *Gypsy12* |  | tgaagacatcatagcagttaccgta | cataattcctcctaacagcacgaa | 1.99 |
| *R4* |  | accaaatctccttattataaatatgagcc | gtgtgtatttggaacagcaatatctatg | 1.97 |
| *Copia1* |  | acttttggctactgaaattggtga | ctttccatgttcaaatgacacgtt | 1.98 |
| *Copia2* |  | cgatacggctaaagaaattctcgaca | ctgatgaaactcctcgtgtataagca | 1.85 |

**Table S2** Relative expression of TGIP marker genes in parental (P_0_) and offspring (F_1_) generation compared to expression in RNAi control* bacterial exposure control group (normalized over two housekeeping genes (n=15). Statistical analysis was performed using Pair Wise Fixed Reallocation Randomization Test (REST2009 software, Pfaffl et al., 2002)

| Gene | Treatment combination | | Paternal generation | | | | Offspring generation | | |
| --- | --- | --- | --- | --- | --- | --- | --- | --- | --- |
|  | RNAi | Heat-killed bacterial exposure | rel. expression | 95% C.I. | p value | rel. expression | | 95% C.I. | p value |
| *hsp83* | *Dnmt2* | bacterial | 0.819 | 0.03 - 9.1 | 0.602 | 1.02 | | 0.49 - 2.11 | 0.846 |
|  |  | control | 1.039 | 0.02 - 8.45 | 0.928 | 1.025 | | 0.39 - 1.97 | 0.815 |
|  |  | naive | 1.414 | 0.12 - 9.88 | 0.272 | 1.036 | | 0.58 - 2.07 | 0.711 |
|  | control | bacterial | 0.885 | 0.11 - 8.42 | 0.7 | 1.059 | | 0.5 - 2.23 | 0.601 |
|  |  | naive | 1.369 | 0.1 - 9.8 | 0.319 | 1.141 | | 0.59 - 2.35 | 0.194 |
| *nimB* | *Dnmt2* | bacterial | 0.938 | 0.001 - 539 | 0.926 | 1.143 | | 0.5 - 5.13 | 0.439 |
|  |  | control | 2.165 | 0.21 - 569 | 0.197 | 1.241 | | 0.63 - 5.34 | 0.16 |
|  |  | naive | 1.23 | 0.01 - 420 | 0.802 | 1.098 | | 0.48 - 4.98 | 0.573 |
|  | control | bacterial | 0.76 | 0.002 - 583 | 0.712 | 1.086 | | 0.58 - 4.52 | 0.578 |
|  |  | naive | 1.17 | 0.01 - 381 | 0.86 | 1.097 | | 0.58 - 4.72 | 0.551 |
| *PGRP* | *Dnmt2* | bacterial | 1.172 | 0.04 - 61.8 | 0.717 | 0.865 | | 0.12 - 14 | 0.627 |
|  |  | control | 1.57 | 0.05 - 79.5 | 0.314 | 1.697 | | 0.13 - 11.5 | 0.583 |
|  |  | naive | 0.816 | 0.03 - 27.2 | 0.605 | 0.979 | | 0.19 - 14.6 | 0.943 |
|  | control | bacterial | 1.136 | 0.03 - 37.8 | 0.75 | 0.769 | | 0.14 - 11.7 | 0.352 |
|  |  | naive | 0.713 | 0.02 - 28.2 | 0.413 | 0.834 | | 0.22 - 9.79 | 0.505 |

**Table S3** Relative expression of TEs after RNAi and heat-killed bacterial exposure treatment in parental (P_0_) generation (expression relative to two housekeeping genes and RNAi control* naïve group, n=6-8). Statistical analysis was performed using Pair Wise Fixed Reallocation Randomization Test (REST2009 software, Pfaffl et al., 2002)

| Gene | RNAi | Heat-killed bacterial exposure | Relative expression | 95% C.I. | p value |
| --- | --- | --- | --- | --- | --- |
|  | *Dnmt2* | bacterial | 0.788 | 0.108 - 4.698 | 0.655 |
| *Gypsy3* |  | naive | 1.078 | 0.363 - 4.200 | 0.78 |
|  | control | bacterial | 0.745 | 0.248 - 3.768 | 0.35 |
|  | *Dnmt2* | bacterial | 1.116 | 0.135 - 8.356 | 0.82 |
| *Gypsy5* |  | naive | 1.189 | 0.202 - 6.986 | 0.682 |
|  | control | bacterial | 0.875 | 0.151 - 6.101 | 0.739 |
|  | *Dnmt2* | bacterial | 0.823 | 0.405 - 1.782 | 0.34 |
| *Gypsy9* |  | naive | 1.053 | 0.621 - 1.996 | 0.733 |
|  | control | bacterial | 0.885 | 0.411 - 2.048 | 0.545 |
|  | *Dnmt2* | bacterial | 0.914 | 0.264 - 4.357 | 0.797 |
| *Gypsy12* |  | naive | 1.074 | 0.415 - 3.917 | 0.779 |
|  | control | bacterial | 0.76 | 0.225 - 3.227 | 0.368 |
|  | *Dnmt2* | bacterial | 0.989 | 0.120 - 2.933 | 0.987 |
| *R4* |  | naive | 1.1 | 0.408 - 2.202 | 0.665 |
|  | control | bacterial | 0.788 | 0.250 - 2.164 | 0.339 |
|  | *Dnmt2* | bacterial | 0.977 | 0.316 - 3.018 | 0.939 |
| *Copia1* |  | naive | 0.92 | 0.273 - 3.261 | 0.786 |
|  | control | bacterial | 0.725 | 0.291 - 2.097 | 0.182 |
|  | *Dnmt2* | bacterial | 0.804 | 0.363 - 1.952 | 0.371 |
| *Copia2* |  | naive | 0.96 | 0.273 - 3.700 | 0.884 |
|  | control | bacterial | 0.696 | 0.226 - 3.045 | 0.235 |

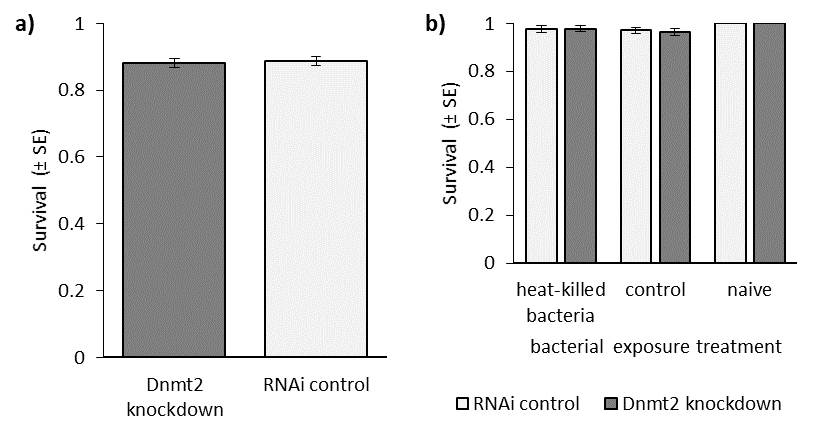

**Figure S1** Survival of RNAi treatment **a)** Proportion of surviving mature adult males seven days post eclosion (n= 8-11 pools of 13-20 individuals per RNAi treatment and replicate). **b)** Proportion of surviving mature adults eight days post eclosion and 24 h post heat-killed bacterial exposure injection (± SEM for three experimental replicates, n=25-70 per RNAi treatment, heat-killed bacterial exposure treatment, and replicate). Different letters indicate significant differences.

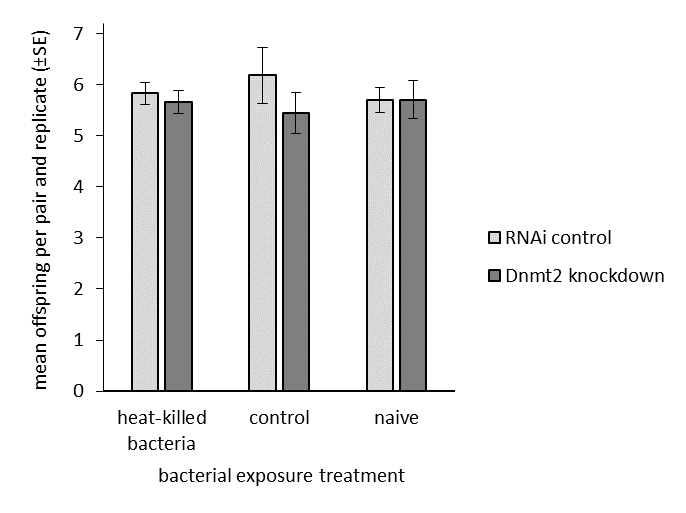

**Figure S2** Male fertility after pupal RNAi treatment and adult heat-killed bacterial exposure injections. Shown is the offspring number from a 24h mating/oviposition period for three replicates (± SE) consisting of 12-50 mating pairs each.

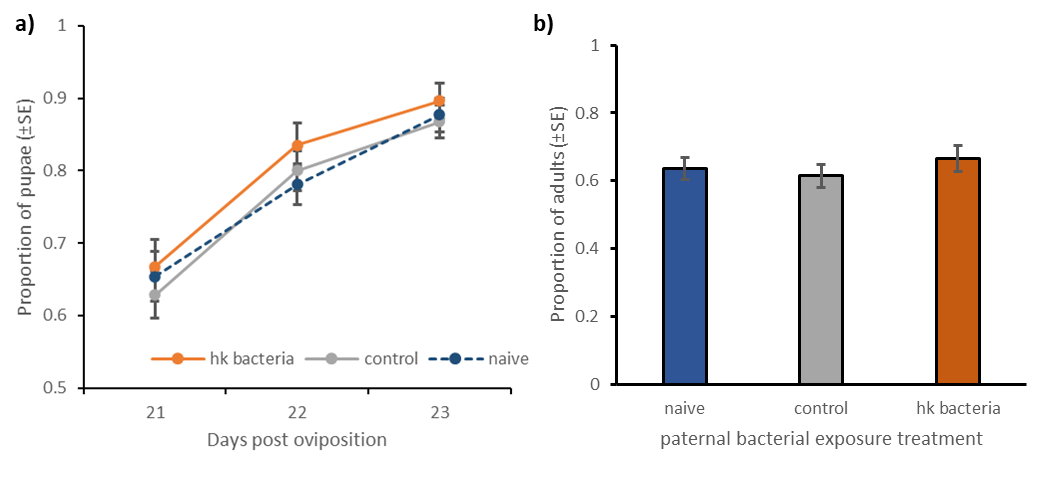

**Figure S3** Development of offspring after paternal heat-killed (hk) bacterial exposure with *B. thuringiensis* **a)** pupation rate (±SE for three experimental replicates) **b)** proportion of eclosed adults (±SE for three experimental replicates) 26 days post oviposition.

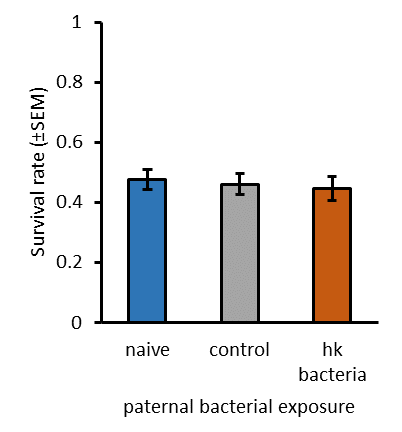

**Figure S4** Survival of F_1_ generation after bacterial challenge according to paternal heat-killed (hk) bacterial exposure treatment. Shown are the proportions of adult offspring that were alive four days post injection with live B. thuringiensis for three independent experimental replicates.

1.         Pfaffl, M. W., Horgan, G. W. & Dempfle, L. Relative expression software tool (REST) for group-wise comparison and statistical analysis of relative expression results in real-time PCR. *Nucleic acids research* **30**, e36 (2002).
